# Integrated epigenome, exome and transcriptome analyses reveal molecular subtypes and homeotic transformation in uterine fibroids

**DOI:** 10.1101/452342

**Authors:** Jitu W. George, Huihui Fan, Benjamin K. Johnson, Anindita Chatterjee, Amanda L. Patterson, Julie Koeman, Marie Adams, Zachary B. Madaj, David W. Chesla, Erica E. Marsh, Timothy J. Triche, Hui Shen, Jose M. Teixeira

## Abstract

Uterine fibroids are benign myometrial smooth muscle tumors of unknown etiology that when symptomatic are the most common indication for hysterectomy in the USA. We conducted an integrated analysis of fibroids and adjacent normal myometria by whole exome sequencing, Infinium MethylationEPIC array, and RNA-sequencing. Unsupervised clustering by DNA methylation segregated normal myometria from fibroids, and further separated the fibroids into subtypes marked by *MED12* mutation, *HMGA2* activation (*HMGA2*hi) and *HMGA1* activation (*HMGA1*hi). Upregulation of *HMGA2* expression in *HMGA2*hi fibroids did not always appear to be dependent on translocation, as has been historically described, and was associated with hypomethylation in the *HMGA2* gene body. Furthermore, we found that expression of *HOXA13* was highly upregulated in fibroids and that overexpression of *HOXA13* in a myometrial cell line induced expression of genes classically associated with uterine fibroids. Transcriptome analyses of the most differentially expressed genes between cervix and myometrium also showed that uterine fibroids and normal cervix clustered together and apart from normal myometria. Together, our integrated analysis shows a role for epigenetic modification in fibroid biology and strongly suggests that homeotic transformation of myometrium cells to a more cervical phenotype is important for the etiology of the disease.

## INTRODUCTION

Uterine fibroids (also known as leiomyomas) are benign tumors that develop in the smooth muscle of the uterine myometrium and are estimated to occur in up to 75% of reproductive age women. While mostly asymptomatic, approximately 25% of women with fibroids suffer from clinically significant symptoms, including pelvic discomfort, menstrual bleeding, menorrhagia to preterm labor, recurrent pregnancy loss and infertility (1,2). There is a strong racial disparity in the disease, with a lifetime prevalence estimated to be 3 times higher in women of African descent (3), who also have earlier clinical onset, higher tumor burden, and greater severity of symptoms. Non-surgical, hormone-based therapies for fibroids offer only short-term mitigation of symptoms, and their use is limited due to significant associated side-effects. Surgical intervention is often the last resort for women seeking permanent relief from the disease. Symptomatic fibroids are the most common indication for hysterectomy in the United States (4).

Approximately 30-40% of fibroids have been reported as having karyotypic abnormalities, the most commonly reported of which are translocations at chromosome regions of 12q15 and 6q21, leading to over-expression of high mobility group AT-hook genes, *HMGA2* and *HMGA1,* respectively (5,6). The HMGA family of proteins are nonhistone, chromatin-binding proteins that regulate transcription by influencing DNA conformation and in the process, accessibility of DNA binding proteins. Due to their various DNA binding properties, they influence diverse cellular processes including cell growth, proliferation, and cell death (7,8).

Whole-exome approaches have recently identified a somatic mutation that is found largely in exon 2 of the Mediator complex subunit 12 (*MED12*) gene and occurs in around 50-70% of the fibroids (9). *MED12* is located on the X chromosome and encodes a highly conserved 250 kDa protein that forms part of the Mediator RNA polymerase II pre-initiation complex. Together, *MED12* mutation (*MED12mt*) and *HMGA1 or HMGA2* overexpression (*HMGA1*hi and *HMGA2*hi, respectively) encompass approximately 80-90% of genetic alterations present in all fibroids (10,11). However, the precise mechanisms disrupted during fibroid development or progression have yet to be determined.

Subtype classification of fibroids based on their mutation status or gene expression characteristics have been proposed (12), but the DNA methylation profiles of these fibroid subtypes have not been reported. Additionally, in some cases, the subtyping was performed without consideration of fibroids from African American women (13). Methylation of cytosine nucleotides 5’ to a guanine (CpG) in DNA is among the most well-established epigenetic marks known to influence gene expression (12), but how these epigenetic modifications might affect transcriptional activity in fibroids has not been well described nor has cytosine methylation in a non-CpG context (CpH methylation) (14,15). The major goal of this study was to delineate the molecular landscape of fibroids based on integrated genome-wide analysis of DNA methylation and mRNA transcription within the context of their mutational status for subtype categorization and identify possible targetable mechanisms for therapeutic intervention.

## RESULTS

### Fibroid Subtype Determination

We applied an integrated approach to study uterine fibroid subtypes by combining DNA methylation array hybridization, whole exome sequencing (exome-seq), and RNA sequencing (RNA-seq) to determine driver mechanisms underlying subtype determination. Ten normal myometrial (5 each Caucasian and African American) and 24 fibroid (12 each Caucasian and African American) samples were collected for DNA methylation analyses. Methylomes for normal myometria and fibroids were profiled using the Infinium MethylationEPIC array (EPIC) (16). Epidemiological studies have well documented a strong racial disparity in the disease, with African-American women presenting with greater incidence, age of onset, and severity (17). Self-identified race for the African American samples was confirmed using EPIC SNP probes (Fig S1A and B) in a constructive predictive model (16) to mitigate confounding effects from possible misidentification because of admixture. To assess cellular composition of the samples, promoter methylation of miR200c/miR141, which are methylated in mesenchymal cells (18) but unmethylated in epithelial cells, was analyzed (Fig S1C). The methylation beta values, corresponding to the fraction of methylated probe signals, suggested very low contamination in the normal myometrium, and approximately 80-90% smooth muscle cells in the fibroids. Further examination of αSMA promoter methylation, which is mostly unmethylated in myofibroblast cells (19), showed consistent results (Fig S1C). Similarly, flow cytometry analysis of human myometrial and fibroid tissue identified approximately 70% and 90% αSMA-positive smooth muscle cells, ensuring a relatively homogeneous population of cells without the need for any further enrichment (Fig S1D). This indicates that our methylation study is a good reflection of the CpG landscape of smooth muscle cells and are not unduly influenced by contaminating cells. Origins of fibroid and normal myometrial samples from specific patients were confirmed by SNP analysis that identified distinct branches in paired normal myometrium and fibroids, while isolated branches were restricted to the unpaired samples (Fig S1E).

Unsupervised clustering of DNA methylation data (Fig 1A) of the most variable 1% CpG sites (Table S1) revealed segregation by disease status (normal and fibroid). Fibroids were further split into three major clades in the dendrogram. Whereas, the *HMGA1*hi fibroids clustered closer to the normal myometrial samples, the *MED12*mt and *HMGA2*hi fibroids clustered closer to each other. Consensus clustering analysis (20) with 1000 iterations showed that the three discovered methylation clusters of fibroids were robust as demonstrated by both sample-based and cluster-based stability scores (Fig 1B). One of the *MED12*mt fibroids showed a high tendency to be clustered with *HMGA2*hi fibroids but predominantly clustered with other *MED12*mt fibroids.

**Figure 1.**
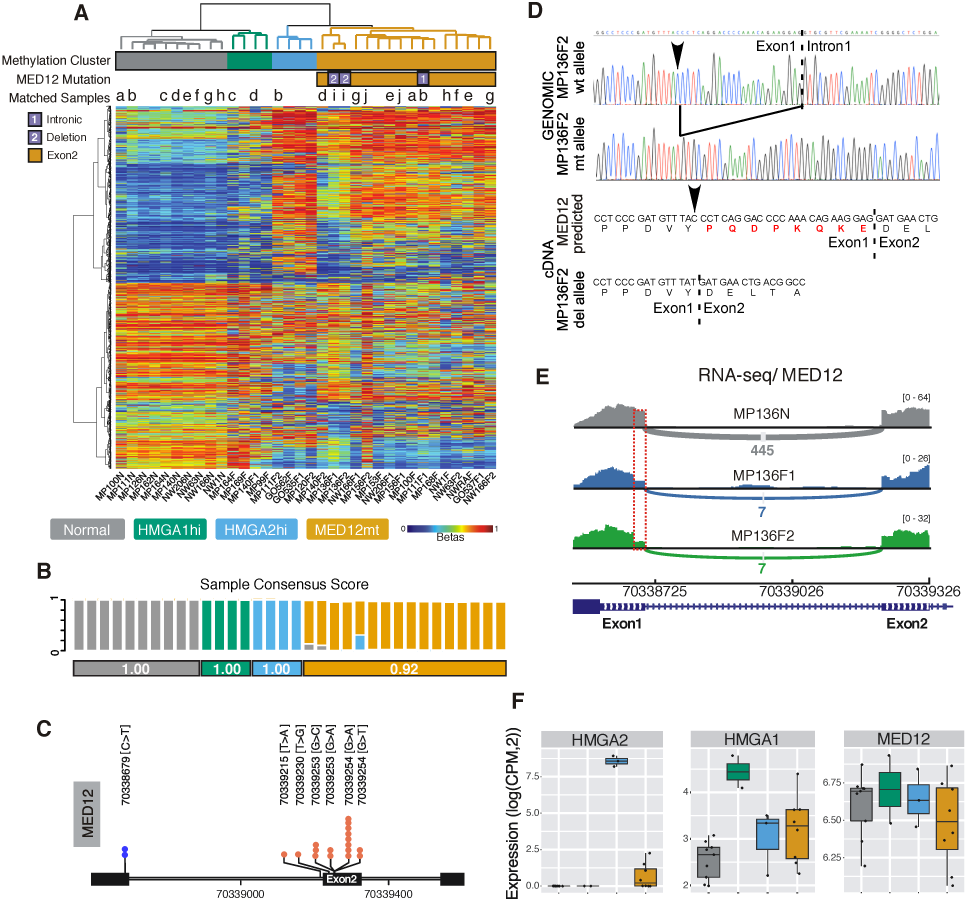
Subtyping of uterine fibroids based on methylome profiling. **(A)** Hierarchical clustering of 10 normal and 24 fibroid samples (columns) on 10,000 most variable CpG sites (rows). A color gradient of blue to red in the heatmap indicates low to high level of methylation (beta values of 0 to 1, corresponding to 0% to 100% methylation). Samples aggregated into four clusters comprising normal myometria, and *MED12*mt, *HMGA2*hi and *HMGA1*hi fibroid subtypes. Mutation status of *MED12* is indicated with a column-side bar, with yellow indicating canonical Exon 2 mutation. Two non-canonical mutations are indicated by numbers. Multiple samples from the same individual are annotated with the same letter at the top of the heatmap. **(B)** Sample-and cluster-stability scores were determined and shown. **(C)** Lollipop plot showing the distribution of different somatic mutations of *MED12* from exome-seq results and Sanger sequencing of PCR products. **(D)** Sanger sequencing electropherogram confirming the novel 24 bp deletion of *MED12* identified from exome-seq data and the C>T mutation (arrowhead). Two genomic clones from MP136F2 are shown. Sequence of mutated cDNA is from amplified PCR product of MP136F2. **(E)** RNA-seq pileup plot for *MED12* exon 1 and exon 2 illustrating decreased reads at the 24-bp deletion (dotted red box). **(F)** Boxplots (boxes, 25–75%; whiskers, 10–90%; lines, median) showing mRNA expression for subtype genes (*HMGA1*, *HMGA2*, *MED12*) for each DNA methylation based cluster/subtype identified in **A**.

We analyzed the fibroids for *MED12* mutation status by Sanger sequencing, and the hotspot, exon 2 mutations in *MED12* (9) were detected in most of the cases (Fig 1C). However, an unreported C>T mutation and a 24 bp deletion from two separate fibroids collected from the same patient (MP136) were also detected using exome-seq (Fig 1D). RNA-seq also showed a clear drop in *MED12* reads in the deleted region of these two fibroids, compared to the rest of the first exon, and to the same region in their matched normal myometrial sample (Fig 1E). Further analysis of the cDNA from these fibroids confirmed the C>T mutation and deletion. Splicing of intron 1 did not appear to have been affected, but the observed in-frame deletion did result in an mRNA with a predicted translated protein missing 8 amino acids (Fig 1D). *HMGA1* and *HMGA2* over-expression marked the two other groups in the non-*MED12*mt fibroids (Fig 1F). *MED12* expression levels were not significantly different between normal myometrium and fibroid subtypes (Fig 1F).

Multidimensional scaling of the whole exome sequencing results showed tight clustering of the myometria and fibroids by patient, confirming that these were matched samples (Fig S2A). We also confirmed that the mutant *MED12* allele was the expressed allele in a few *MED12mt* fibroids (Fig S2B). With both exome sequencing and RNA sequencing data, we were able to investigate whether the paired fibroids from the same patients (MP111 and MP136) came from a single cell or had separate origins by examining X chromosome inactivation patterns. One of the two X chromosomes is randomly inactivated early in development, and fibroids with different inactive X chromosome would be unlikely to come from the same cell of origin. MP111F1 and F2 fibroids expressed alternative alleles for heterozygous loci on chromosome X (Fig S2C), as well as poorly correlated methylation patterns (see Fig S2D, also Fig 1A and Fig S4B), suggesting that they originated independently from two cells with a different chromosome X inactivated (Fig S2E). In contrast, MP136F1 and F2 expressed the same alleles (Fig S2F), as well as highly similar methylation patterns (see Fig S2G, also Fig 1A and Fig S4B), suggesting a probable single cell of origin (Fig S2H). Conflicting results regarding clonality have been reported; however, previous exome sequencing of fibroids (11) and these results suggest that clonality can vary among the fibroids.

Mutational burden of fibroids was generally less than 0.5/MB (Fig S3A), except for MP164F (2.5/MB), and comparable to that of pediatric leukemias and lymphomas, which represent some of the lowest in human cancers profiled to date (21). A prevalence for C>A mutation in some of the fibroids (Fig S3B) was observed, different from the common C>T mutation in CpG context. However, association of the fibroids with mutation signatures did not reveal any overt differences within and between the *MED12*mt and *HMGA2*hi fibroid subtypes (Fig S3C), neither were we able to identify complex chromosomal rearrangements in our *MED12*mt samples by available bioinformatic tools suggesting that mutational burden is not a major contributor to the fibroid phenotype in our samples.

### DNA Methylation Landscape Is Altered in Fibroids

Overall distribution of CpG and CpH (or non-CpG) methylation were largely unremarkable in all samples (Fig S4A). At loci with fibroid-specific hypermethylation compared to normal myometria, *HMGA1*hi fibroids were closest to normal myometrial samples (Fig S4B), similar to the clustering results with the top 1% most variably methylated sites (Fig 1A). *HMGA2hi* and *MED12*mt fibroids had elevated levels of methylation at CpH sites (Fig S4C). *HMGA2*hi had the highest-level gain of methylation at CpG and CpH sites among all groups, which is consistent with the observation that they also had the highest expression level of the *de novo* DNA methyltransferase DNMT3A (Fig S4D).

We examined the sites interrogated on this array by various genomic compartments. CpG islands were largely unmethylated in both normal myometria and fibroids, and highly methylated domains (HMDs) in myometria remained highly methylated in fibroids (Fig S4E). We also did not observe significant hypomethylation in the partially methylated domains (PMDs), even though loss of methylation within PMDs, particularly in the context of WCGW (where W=A or T) without neighboring CpGs (dubbed solo-WCGW), has been suggested to track accumulation of cell divisions in normal cells and is commonly observed in tissues that have undergone extensive clonal expansion like cancer (22). Methylation at enhancer regions, however, exhibited a small shift in the overall distribution between normal myometria and fibroids from PMDs to HMDs (Fig S4E).

Enhancer activity landscape can be inferred from DNA methylation profiles (23), with unmethylated distal regions usually marking active enhancers. A large fraction of the probes on the EPIC array interrogate distal elements (defined as +/− 2kb away from the TSS) containing at least one binding site for each of 158 transcription factors (TFs) that we previously annotated (16), based on ENCODE ChIP-seq data (24). We assessed enrichment or depletion of differentially methylated cytosines (DMCs) in binding sites for the TFs by hypergeometric testing (FDR =1*10-6), comparing all fibroid and each of the fibroid subtypes to normal myometria (Fig S5). *HMGA1*hi fibroids were similar to normal myometria and had few DMCs. *MED12*mt and *HMGA2*hi fibroids exhibited similarity in TFBS enrichment at their hypermethylated distal loci, further indicating that they could share similar transcriptional rewiring. Notably, binding sites for EZH2 and SUZ12, components of the PRC2 complex (25), were highly enriched in the hypermethylated, and likely closed off, cytosines. In contrast, estrogen receptor-α (ERα) binding sites were enriched in distal sites that lose methylation and are presumably activated in the fibroids.

### DNA Hypomethylation in the HMGA2 Gene Body Correlates with Higher HMGA2 Expression

Compared to normal myometria, two adjacent CpG sites in a CpG island within the *HMGA1* promoter gained DNA methylation in *MED12*mt and *HMGA2*hi fibroids, but remained hypomethylated in *HMGA1*hi fibroids, in line with the high expression level of *HMGA1* observed in *HMGA1*hi fibroids (Fig S6; Fig 1). In contrast, a segment of the gene body of *HMGA2* (measured by 13 consecutive DNA methylation probes) was hypomethylated in *HMGA2*hi fibroids, compared with other fibroids and to normal myometria (Fig 2A). Upregulation of *HMGA2* expression has been generally attributed to rearrangements at 12q14-15 (26,27). However, when fluorescent *in situ* hybridization (FISH) was performed on fresh-frozen tumor sections or exponentially growing fibroid cell cultures of samples identified as *HMGA2*hi, 12q14-15 rearrangement was not detected in either GO535F1 or MP120F2 with probes over 600 KB upstream of *HMGA2* (Fig S7A and B). qRT-PCR was performed to confirm the assignment of these fibroids to the *HMGA2*hi subtype by the RNA-seq results (Fig S7C). The 3’ end distal hypomethylated CpG site was located within a binding motif for CTCF, as determined by ENCODE ChIP-seq data (Fig 2A) (28). CTCF is involved in forming long range chromatin loops that alter the 3D structure of chromosomes and acts as an insulator of transcriptional activity of the encompassed genes (29). To locate CTCF binding regions upstream of, and within the *HMGA2* gene body, we analyzed enhancer-promoter interactions from FANTOM5 (black lines), and overlaid it with available ChIA-PET interaction data (red line). We also inferred open and closed compartments (A/B compartment) (30) using DNA methylation profiles surrounding the *HMGA2* region (Fig 2B). Indeed, the chromatin was open for this locus in the *HMGA2*hi subtype specifically and closed in the others. These results suggest that epigenetic alteration could be an additional mechanism allowing for overexpression of *HMGA2,* at least in some fibroids.

**Figure 2.**
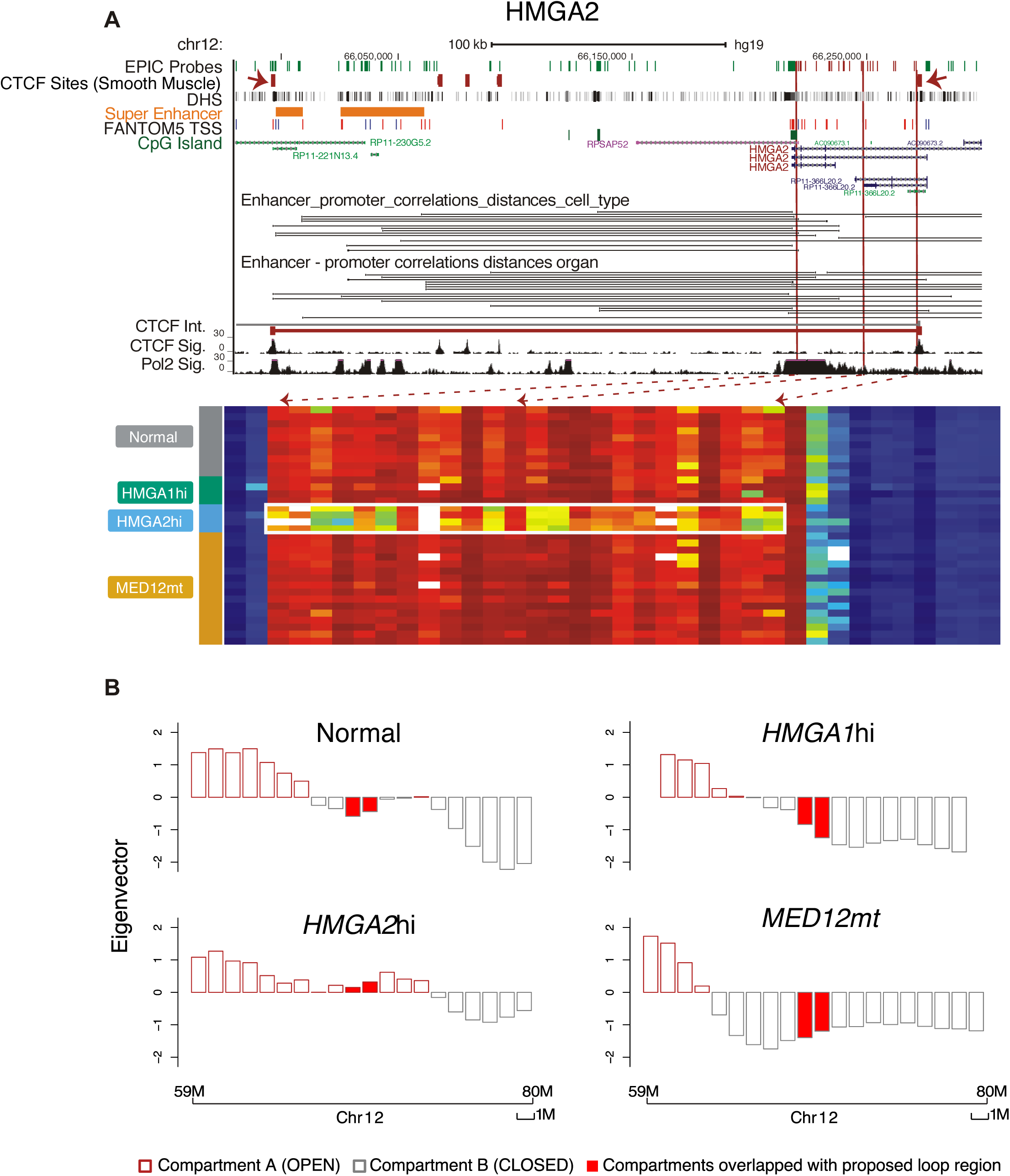
*HMGA2* gene body hypomethylation and altered chromatin organization in *HMGA2* hi fibroids. **(A)** UCSC genome browser view of gene *HMGA2*, coupled with DNA methylation level for this region shown as a heatmap (columns ‐ CpGs; rows ‐samples grouped by subtype). Top tracks indicate locations of EPIC probes, smooth muscle CTCF ChIP-seq peak, DNase I hypersensitivity sites (DHS), super enhancer, TSS, CpG island, UCSC gene, enhancer-promoter correlations, CTCF looping. CTCF signal and Pol2 signal are added on top of the gene heatmap. White box in the heatmap highlights the location of gene body hypomethylation in the *HMGA2* subtype. Red dashed arrows indicate the corresponding genomic locations of those hypo-methylated probes. Two red solid arrows in the CTCF track show the location of CTCF in smooth muscle cells, in line with predicted CTCF interaction loop boundaries curated from ChIA-PET. **(B)** Reconstruction of A/B compartments using EPIC array data on chromosome 12 in each of the four DNA methylation groups. The CTCF loop region identified in 2A is indicated with filled red bars, which switches from B to A compartment specifically in the *HMGA2*hi subtype.

### Transcriptome Analyses Identifies Multiple Commonalities and Differences Between Fibroid Subtypes

As with DNA methylation, global RNA-seq analyses showed that, in addition to clustering separately from normal myometrial samples, the *MED12*mt, *HMGA1*hi, and *HMGA2*hi subtypes also clustered separately from each other and independently of patient origin (Fig 3A). Gene set enrichment analyses of the RNA-seq results between the *MED12*mt or *HMGA2*hi fibroids, when each was compared with normal myometria, showed a large number of shared activated and repressed genes among the top-ranked gene sets (Fig 3B). More than half of the up-regulated genes in *HMGA2*hi fibroids were similarly regulated in the *MED12*mt fibroids (Fig 3C), and nearly half of the down-regulated genes in *HMGA2*hi fibroids were also down regulated in *MED12*mt fibroids (Fig 3D). Gene ontology analyses of the differentially expressed genes showed a high concordance of dysregulated genes (Fig 3E). KEGG pathway analyses of the differentially expressed genes showed that a few of the pathway changes were shared between *MED12*mt and *HMGA2*hi fibroid subtypes. Concordant with previous transcriptomic profiling, we identified elevated expression of *RAD51B*, *PLAG1* and *PAPPA2* in our *MED12*mt, *HMGA2*hi and *HMGA1*hi fibroids, respectively (Table S2) (13). These RNA-seq results suggest that the *MED12*mt and *HMGA2*hi fibroid subtypes are more alike transcriptomically than they are different.

**Figure 3.**
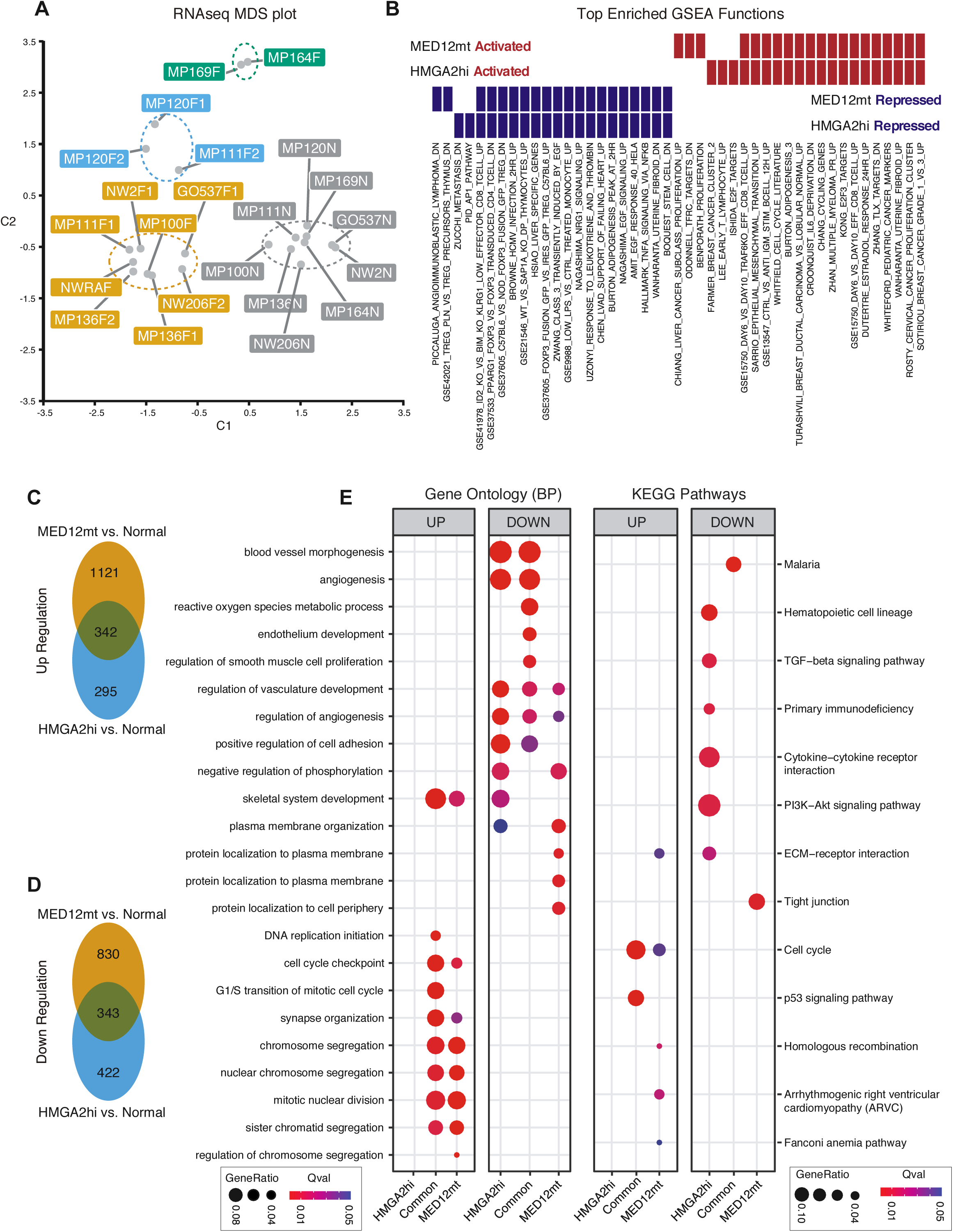
Transcriptome characterization of the three fibroid subtypes. **(A)** MDS plot based on RNA-seq gene expression data. Each dot represents one sample, colored by DNA methylation-based subtypes. **(B)** GSEA analysis of up‐ or down-regulated genes in *MED12*mt and *HMGA2*hi subtypes compared to normal samples. The topmost 20 enriched gene sets for genes upregulated (red) or down regulated (blue) in each fibroid subtype versus normal myometrium comparison are shown. Venn diagram illustrates the overlap for up-regulated **(C)** and down-regulated genes **(D)** between *MED12*mt and *HMGA2*hi subtypes. **(E)** Gene ontology (biological process) and KEGG pathways analysis for *HMGA2*hi-specific, shared, and *MED12*mt-specific up-regulated (left) and down-regulated (right) gene sets. Gene enrichment ratio and significance level are shown by the size and color of each circle, respectively.

In Fig 1F, we showed that the average *HMGA2* expression level was also significantly elevated in *MED12*mt fibroids (log 2 FC=3.2), though to a lesser degree than in *HMGA2*hi fibroids (Log2 FC=11.6). Further analysis of the RNA-seq results identified that, although most of the *MED12*mt fibroids did not express any *HMGA2* transcripts, 3 *MED12*mt fibroids did express *HMGA2* (Fig S8), suggesting that there might be two subtypes of *MED12*mt fibroids, *MED12*mt and *MED12*mt/*HMGA2* expressing, with different transcriptional profiles. However, these *MED12*mt/*HMGA2* expressing fibroids did not cluster closer to the *HMGA2hi* fibroids in either the RNA-seq (Fig 3A) or DNA methylation heatmaps (Fig 1A).

To identify genes dysregulated due to altered promoter methylation level, we integrated DNA methylation and RNA-seq profiles for each fibroid subtype (Fig 4). Genes with significantly altered promoter CpG methylation (absolute delta β values>0.25; p<0.05) and associated gene expression change between normal myometrium and fibroids are listed in Table S3. A few hypomethylated and induced genes were identified, but most (*VCAN*, *RAD51B*, *COL1A1*, etc.) have been previously reported. *KRT19*, which has also been previously reported to be silenced by promoter hypermethylation in fibroids (31), was the most down-regulated gene in all fibroids compared normal myometria using this analysis (Fig 4A). The heatmap of beta values for the EPIC probes in the *KRT19* gene show that most of the hypermethylation in the fibroids is occurring in the promoter region (Fig 4B). We also identified many additional genes similarly silenced by promoter methylation, including genes involved in the retinoic acid pathway (*ADH1B*), WNT pathway (*WNT2B*), and stem cell function (*GATA2*, *KLF4*) (Fig 4A). Many of these are known tumor suppressor genes, silencing of which could be important for fibroid growth. More granular analysis of *KLF4* showed that it was hypermethylated and downregulated in each of the fibroid subtypes compared to normal myometria (Fig 4C). We then analyzed coordinated differential methylation and gene expression by fibroid subtypes. In *HMGA1*hi fibroids *SMOC2* as the most hypermethylation and down regulation gene (Fig 4D). SMOC2 can stimulate endothelial cell proliferation and migration, which is consistent with the observed hypoxia in fibroids (32,33). *MED12*mt fibroids (Fig 4E) had a differential gene pattern similar to that of total fibroids. Several of the differentially regulated genes (*PAPPA2*, *PLAG1,* for example) in *HMGA2*hi fibroids have also been previously described. However, *HOXA13* was identified to be hypomethylated and upregulated in *HMGA2*hi fibroids compared to normal myometria (Fig 4F). Given the importance of this HOX gene in female reproductive tract development, we continued to further investigate the HOXA loci.

**Figure 4.**
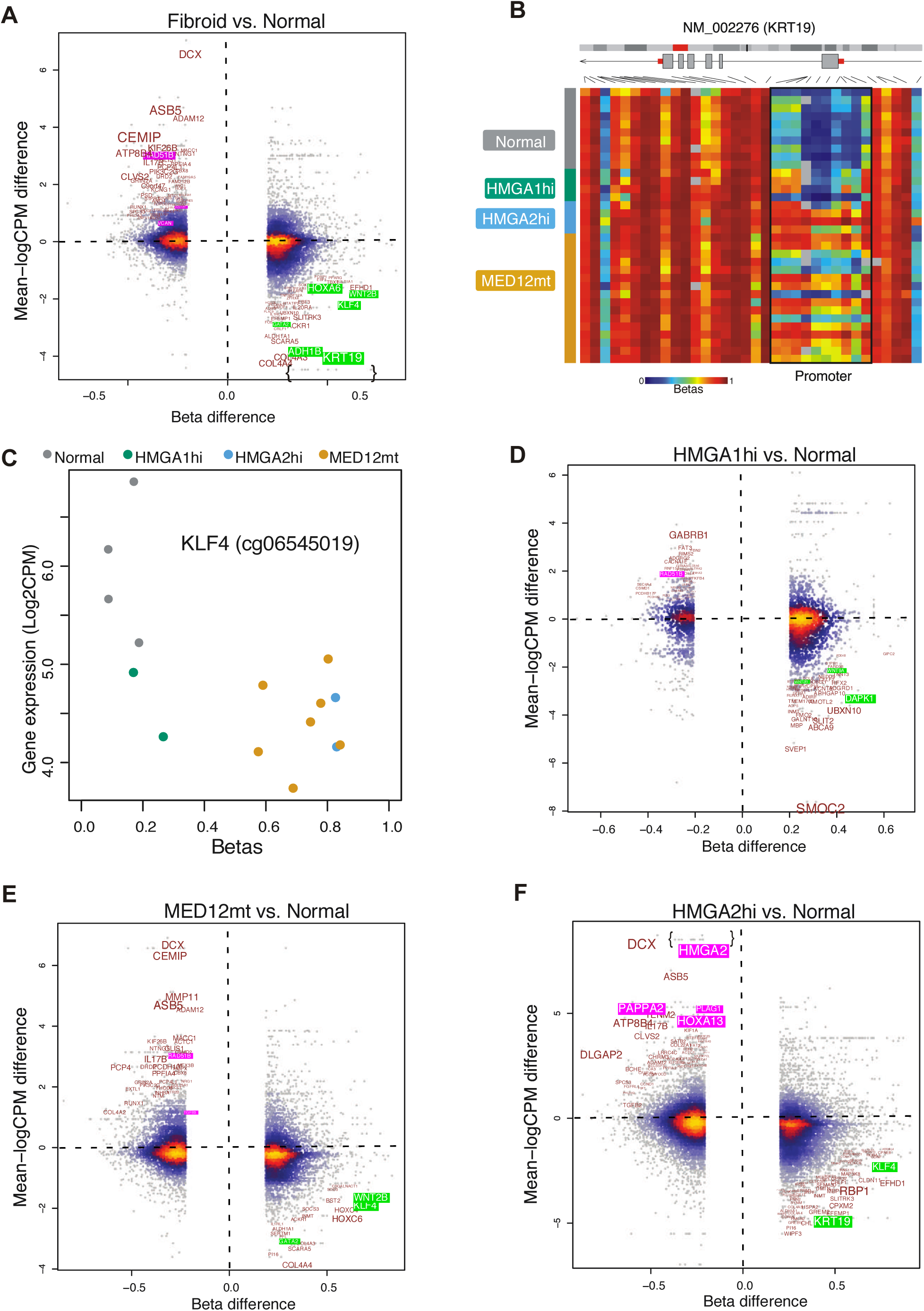
Differential promoter DNA methylation versus differential gene expression. Promoter DNA methylation beta difference is plotted on the x-axis, with log2 fold change for the corresponding gene is plotted on the y-axis. This analysis is done for all fibroids **(A),** *HMGA1*hi **(D),** *HMGA2*hi **(E)**, and *MED12*mt **(F)** versus normal samples. Color represents local dot density. Green-highlighted genes are some of the most correlated between hypermethylation and its down-regulation of expression. Pink-highlighted genes are some of the most correlated with hypomethylation and induced expression. Text is sized to account for both probe methylation difference and corresponding gene fold change. **(B)** *KRT19*, a representative epigenetically silenced gene, is further visualized in detail as a heatmap with multiple probes called within the same gene. **(C)** Dotplot (probe methylation is plotted as x axis and related gene expression as y axis) of KLF4 showing down-regulation of gene expression is observed in all fibroid subtype.

### HOXA13 Induces a Homeotic Transformation in Myometrium

The HOX family of homeobox genes dictates normal development of the urogenital tract (34-37) and their expression is regulated by direct action of estrogen and progesterone (38). Interestingly, the fibroids clearly exhibited a switch to expression of more posterior HOXA genes (Fig 5A), based on RNAseq data. Among the HOXA genes, only *HOXA13* mRNA expression reached statistical significance after correcting for multiple comparisons on a genome-wide survey. It was highly expressed in *MED12*mt (Log2 FC= 3) and *HMGA2*hi (Log2 FC=4.4) fibroids compared with either normal myometria or *HMGA1*hi fibroids (Fig 5B. We confirmed high *HOXA13* mRNA abundance in a validation set of fibroid samples, compared to adjacent normal myometria by qRT-PCR (Fig 5C). The long non-coding RNA (lncRNA), HOXA transcript at the distal end (*HOTTIP*), which is located at the 5’-end of the HOXA cluster and coordinately regulated with that of multiple HOX genes (39), including *HOXA13*, was also elevated (Fig S9A; S9B). qRT‐ PCR revealed a high correlation between *HOXA13* and *HOTTIP* mRNA levels in normal myometrium and uterine fibroids (Fig S9C), suggesting coordinated expression or even a potential interrelated feed-forward mechanism driving uterine fibroid growth.

**Figure 5.**
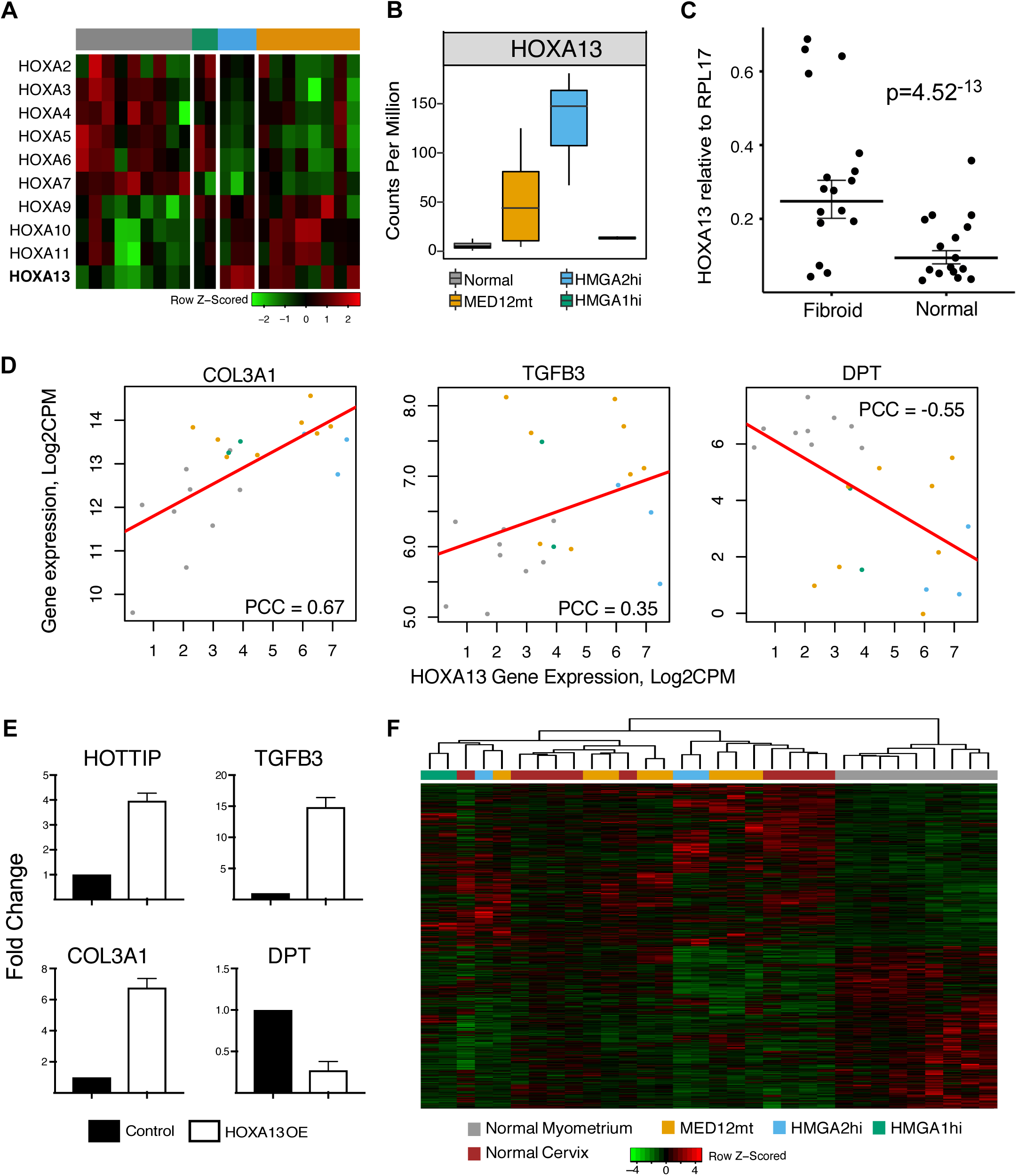
*HOXA13* overexpression in normal myometrium leads to homeotic transformation. **(A)** Boxplots (boxes, 25–75%; whiskers, 10–90%; lines, median) showing *HOXA13* gene expression in normal myometrium and, *MED12*mt, *HMGA2*hi and *HMGA1*hi fibroids from the RNA-seq results. **(B)** Relative expression of *HOXA13* by qRT-PCR compared to RPL17 housekeeping gene in a validation set of samples between normal myometrium (n=17) and fibroids (n=19). **(C)** Expression of *HOXA13* (x axis) versus that of fibroid-characteristic genes (y axis of each panel, for *COL3A1*, *TGFB3*, *DPT*) respectively. Red lines represent linear regression trends, and the Pearson’s Correlation Coefficient (PCC) are indicated for each analysis. **(D)** qRT-PCR analyses of *HOTTIP*, *TGFB3*, *COL3A1*, and *DPT* mRNA expression in control and untransfected UT-TERT cells, compared to UT-TERT cells transfected with HOXA13. Mean fold change normalized to control is shown, with error bars representing SEM. **(E)** Fibroids samples clustered together with normal cervix instead of normal myometrium based on 5,000 most variably expressed genes (rows in heatmap) between normal cervix and normal myometrium.

The expression of a number of smooth muscle cell and extracellular matrix genes are known to be altered in uterine fibroids compared to myometria. *HOXA13* expression in the fibroids positively correlated with the expression levels of *COL3A1* (40) and *TGFB3* (40), and negatively correlated with that of *DPT* (41) (Fig 5C), genes classically associated with the fibroid phenotype. When transfected with a *HOXA13* over expression vector, the UT-TERT myometrial cell line had significantly altered expression of these genes and that of *HOTTIP,* compared to control, untransfected UT-TERT cells (Fig 5D), indicating that HOXA13 can likely regulate the expression of these characteristic fibroid genes.

Developmentally, *HOXA13* expression is posteriorly restricted in the differentiating Müllerian ducts and is required for normal cervix and vagina development (42). Gene expression profiles between normal myometrium and normal cervix (GSE55940 (43) and GSE63678 (44)) were compared to uterine fibroids and identified several genes that are associated with fibroids, including collagens (*COL3A1*, *COL1A1*), matrix metalloproteases (*MMP14*, *MMP16*), and a member of the TGF-β superfamily (*TGFB3*). Clustering analysis based on this differential gene set showed that uterine fibroids clustered more closely to normal cervix than to myometria (Fig 5E), strongly suggesting that the development of uterine fibroids is in fact a homeotic transformation into cervical tissue results from induced expression of *HOXA13* in normal myometrium.

## DISCUSSION

In an attempt to understand the tumorigenic role of *MED12* mutation in fibroid development, a genetically modified mouse model has been developed that shows expression of the *MED12* mutant transgene on either a *MED12* null or WT background leads to fibroid formation, suggesting that *MED12* mutation could drive their development through a gain-of-function or dominant negative mechanism (45). In contrast, biochemical assays demonstrate *MED12* mutations lead to loss of CycC-CDK8/19 binding and function, suggesting that *MED12* mutations might be a loss-of-function phenotype (46). Biochemical analysis further revealed quantitative differences within the various *MED12* exon 2 mutations in regards to kinase activity indicating that maybe all *MED12* mutants are not equal (46). Consideration of exon 1 mutations (9), including the 8-amino acid deletion described here (Fig 1), and the clustering of these fibroids in both DNA methylation and RNA-seq analyses with other *MED12* mutants (Fig 1; Fig 3), suggests that a loss of function is more likely. However, given the hotspot *MED12* mutations in fibroids, and now the in-frame deletion described in Fig 1, further study will be necessary to determine the precise mechanisms disrupted by *MED12* mutation in fibroids.

Over-expression of *HMGA2*, a gene normally not highly expressed in normal myometrium, is the second most common phenomena that is known to occur in fibroids (5). *HMGA2* over-expression has been attributed to genetic alterations involving translocations or aberrant splicing within the coding region (47,48). *HMGA2* expression has also been shown to be potentially regulated by the microRNA Let-7 family (49,50). A gain of function mechanism involving fusion of *RAD51B* to *HMGA2* has also been identified to contribute to overexpression (51), however, we were unable to detect *RAD51B*-*HMGA2* fusion or aberrant splicing in our exome and RNA-seq analyses. Using FISH probes spanning increasing distances upstream of *HMGA2* gene body, translocations were confirmed in only one of the *HMGA2*hi fibroids analyzed. The hypomethylation we observed in all *HMGA2*hi fibroids suggests that epigenetic alteration could be another mechanism involved in *HMGA2* upregulation. This hypomethylated region encompass a binding site for CTCF, a ubiquitously expressed zinc finger protein that has been shown to regulate gene expression by mediating inter and intra-chromosomal interactions between distal genomic sites that are sensitive to DNA methylation (29,52). The identification of CTCF motifs within the *HMGA2* gene body led us to hypothesize that hypomethylation might be associated with altered CTCF-mediated looping which would facilitate interaction between a distal enhancer and the *HMGA2* gene promoter leading to over-expression of *HMGA2*. Biochemical confirmation of CTCF binding and activation of HMGA2 gene expression in uterine fibroids will need to be performed.

Enhancers have gathered increasing research interest as the most dynamically used compartment of the genome (53-55). Previously published genome-scale DNA methylation studies utilized the Illumina Infinium HumanMethylation27 (31) or HumanMethylation450 (HM450) (56) arrays, which largely focused on the promoter regions. The MethylationEPIC array, which superseded the HM450 array, boasts an unparalleled coverage for enhancer sites (16). We were able to infer the epigenetic regulatory landscape of fibroids by analyzing the methylation states of these sites. Binding sites for EZH2 and SUZ12, both components of the Polycomb complex, were most enriched for hypermethylated sites in fibroids compared to normal myometria. Polycomb Group proteins form large heteromeric complexes involved in reversible transcription repression, of which Polycomb repressive complexes 1 and 2 (PRC1 and PRC2), have been characterized most extensively (57) as master regulators of stem cell differentiation. PRC binding sites have been shown to be highly enriched for sites hypermethylated in cancer (58) and presumably maintain a ‘locked-in’ stem-like signature in malignant cells. While recent studies (59) and our study (Table S2) have shown higher expression of EZH2 in uterine fibroids, analyses of altered EZH2 expression and binding and the resulting changes in target gene expression in uterine fibroids to determine mechanisms of action and possible therapeutic potential have yet to be reported. Our analysis also identified the binding sites for NANOG, a transcription factor pivotal for self-renewal and ground state pluripotency of embryonic stem cells (60), were also selectively methylated in both *MED12*mt and *HMGA2*hi fibroids. Consistently, one of the Yamanaka pluripotency inducing factors, *KLF4*, was also epigenetically silenced in fibroids (Fig 4C). We and others have hypothesized that fibroids evolve from myometrial stem cells that have undergone genetic modifications (1,2). Furthermore, lower stem cell population have been reported in uterine fibroids compared to normal myometria (61,62). Loss of KLF4 expression and closed NANOG binding sites observed in this study are both in line with such hypothesis. We also identified hypomethylated DNA methylation in TFBSs of Estrogen Receptor α, ERα, (*ESR1*) in all the fibroids, excluding the *HMGA1*hi subtype, and that of Glucocorticoid Receptor, GR, (*NR3C1*), particularly in *MED12*mt fibroids (Fig S5). A number of studies have reported higher expression of *ESR1* in uterine fibroids compared to normal myometria (63-65), and treatment of cultured fibroid cells with estrogen can increase proliferation and cell cycle progression (66). In contrast, GR, another member of the nuclear steroid receptor superfamily, has been postulated to be antagonistic to the estrogen-induced response in fibroids (67-69). *HMGA2*hi fibroids have been reported to be larger in size compared to *MED12*mt tumors (70,71). Perhaps the enrichment of GR TFBSs among hypo-methylated loci in MED12mt fibroids particularly antagonizes the estrogen-induced effects, such as controlling fibroid size. ChIP-seq and functional assays will need to be performed to determine whether altered methylation at these sites is important for TF binding and whether these TF binding differences impact fibroid biology in a meaningful way.

HOX genes encode a supercluster of homeobox transcription factors that are highly conserved, critical regulators for proper development of the female reproductive tract (38). Along the cranial to caudal axis of the differentiating Mu□llerian duct, *HOXA10* is expressed in the uterus while *HOXA13* is expressed in the cervix and vagina (34,42). *HOXA13* expression has been reported in a number of solid tumors including prostate cancer, gastric cancer, glioblastoma multiforme, ovarian cancer and hepatocellular carcinoma (72-75). Replacement of the *HOXA11* homeobox gene with *HOXA13*, leads to homeotic transformation of uterus towards posterior cervix and vagina (36). This homeotic transformation confirms that *HOXA11* and *HOXA13* are not functionally redundant while strengthening the role of HOX genes in body patterning. Among the HOX genes that we analyzed, we identified *HOXA13* to be uniquely upregulated within the HOX gene family in uterine fibroids, which has not been previously appreciated. Our results strongly suggest that expression of *HOXA13* in uterine fibroids drives aberrant gene expression in myometrial cells, transforming them into uterine fibroids with a gene expression profile that is very characteristic of cervix. Much like fibroids, the cervix, a collagen dense tissue that protects the uterus and, if gravid, the fetus from pathogenic assault, undergoes significant remodeling postpartum. Thus, development and resolution of fibroids could be controlled by mechanisms similar to those observed in the cervix. For example, progesterone can keep the cervix stiff and help prevent cervical softening or ripening (76), and is normally required for fibroid growth and development (77). At term, mechanisms driving cervical ripening, characterized by partial dissolution of the collagen matrix is necessary for delivery (78), could also be driving the decrease in fibroid burden observed in postpartum uteri (79). Relaxin is a peptide hormone associated with cervical ripening at parturition (80), but it’s possible role in fibroid biology needs to be investigated in light of our results showing the transformation of fibroids to a more cervix-like tissue. During parturition, the lower uterine segment of the myometrium undergoes a regionalization event leading to increased contractility, a fundamental phenomenon of spontaneous labor (81). This contractile phenotype has now been attributed to higher expression of *HOXA13*, a key regulator of a number of genes that are involved in cell contractility and cell-cell adhesion, further associating the role of *HOXA13* in myometrial transformation (82).

In conclusion, using an integrative approach of RNA-seq, DNA methylation array and exome sequencing, we characterized the genetic and epigenetic profiles of uterine fibroids, which clearly allows for their subtyping by mutation and altered methylation. Our analysis also identified hypomethylation of *HMGA2* gene body as a possible mode of regulation that is independent of translocation. We also showed high correlation with gene expression and DNA methylation, highlighting the regulatory potential of altered DNA methylation driving development of uterine fibroids. Transcription factor binding analysis further identifies fibroid subtype specific regulators and hint at critical role of these regulators in fibroid tumorigenesis. Finally, a deeper characterization of our RNA-seq results identified expression of a HOX gene, *HOXA13*, deregulation of which influences a number of genes characterized in fibroid biology. Importantly, we identified a homeotic transformation of normal myometrium to cervix-like tissue, probably through the subtype-independent upregulation of *HOXA13* expression. Future work will validate the mechanisms altered by *HOXA13* overexpression in fibroid biology and its possible targeting for therapeutic intervention for symptomatic patients.

## METHODS

### Sample processing

Fibroid and matched myometrial samples were collected from hysterectomies of consented patients using Spectrum Health or Northwestern University IRB approved protocols for secondary use of biobank tissues. Samples were aliquoted upon arrival for DNA and RNA isolation, and cell separation. Race was confirmed essentially as described in (16). *MED12* mutation was determined by PCR amplification followed by Sanger sequencing using primers 5’-CTTCGGGATCTTGAGCTACG-3’ and 5’-GGAGGGTTCCGTGTAGAACA-3’ for Exon1, primers 5’-GCTGGGAATCCTAGTGACCA-3’ and 5’-GGCAAACTCAGCCACTTAGG-3’ targeting Exon 2. MED12 cDNA was amplified using primers 5’-CTTCGGGATCTTGAGCTACG-3’ and 5’-AAGCTGACGTTCTTGGCACT-3’ spanning Exon 1 and Exon 2. HMGA1 AND HMGA2 overexpression was determined by RNA-seq and/or confirmed by qRT-PCR with the following primers 5’-GAAGTGCCAACACCTAAGAGACC-3’ and 5’-GGTTTCCTTCCTGGAGTTGTGG-3’ and 5’-GAAGCCACTGGAGAAAAACGGC-3’ and 5-GGCAGACTCTTGTGAGGATGTC-3’, respectively. Promoter (defined as +/− 500bp of transcription start site (TSS)) methylation status of miR-200 family (83,84) and alpha smooth muscle actin (α-SMA) (85) were also employed to validate the major component of the cells analyzed as smooth muscle.

### Flow Cytometry

Tissues (two matched normal and fibroid samples) were minced, placed in digestion media (DMEM/F12, 1X antibiotic-antimycotic, 10% FBS, 2 mg/ml Collagenase Type I (Sigma), 1 mg/ml DNase Type I (Sigma), and 5 mM MgCl2) and incubated at 37° C overnight with agitation. Cell suspensions were passed through 100 μm and 40 μm cell strainers, washed with PBS, and centrifuged. The cell pellets were resuspended in 5 ml ACK Lysing Buffer (Thermo Fisher Scientific) to remove red blood cells, washed with PBS, centrifuged, and resuspended in 2% PFA for 10 min. Following PBS wash and centrifugation, cells were permeabilized in ice-cold methanol. Cells were washed again then incubated with anti-Actin α-smooth muscle-cy3 (Sigma, cat #C6198, 1:500) in PBS with 0.5% NP-40 for 1.5 h at RT. Cells were washed and resuspended in PBS and analyzed on an Accuri C6 flow cytometer (BD Biosciences, San Jose, CA). Events were gated initially by forward and side scatter, then for singlets (side scatter area × height) and finally for Cyanine 3 (Cy3) fluorescence using FlowJo software (FlowJo, Ashland, OR). Unstained cells served as a gating control.

### Fluorescence in situ hybridization (FISH)

FISH probes were prepared from purified BAC clones from the BACPAC Resource Center (bacpac.chori.org). The BAC clones are as follows; Probe set 1: CH17-111D2 (12q14.3-1 green) and CH17-63I9 (12q14.3-2 orange), Probe set 2: CH17-392C11 (12q14.3-3 green) and CH17-63I9 (12q14.3-2 orange), and Probe set 3: CH17-305B19 (12q14.3-5 green) and CH17-63I9 (12q14.3-2 orange). FISH probe 12q14.3-2 is located at the distal end of gene *HMGA2*, while probes 12q14.3-1, 12q14.3-3, and 12q14.3-5 are all proximal of *HMGA2*. FISH probe 12q14.3-2 was labeled with Orange-dUTP and all other probes were labeled with Green-dUTP (Abbott Molecular Inc., Abbott Park, IL), by nick translation. Tumor touch preparations were prepared by imprinting thawed tumors onto positively-charged glass slides. The sample slides were fixed in methanol:acetic acid (3:1) for 30 min and air-dried. Slides were then aged in 2X saline/sodium citrate (SSC) at 60 °C for 20 min, digested with 0.005% pepsin at 37 °C for 5 min, and washed with 1X PBS for 5 min. Slides were placed in 1% formaldehyde/PBS for 10 min at room temperature, washed with 1X PBS for 5 min, and dehydrated in an ethanol series (70%, 85%, 95%) for 2 min each. Slides were then denatured in 70% formamide/2X SSC at 74 °C for 3.5 min, washed in a cold ethanol series (70%, 85%, 95%) for 2 min each, and air-dried. The FISH probes were denatured at 75 °C for 5 min and held at 37 °C for 10-30 min until 10 ul of probe was applied to each sample slide. Slides were coverslipped and hybridized overnight at 37 °C in the ThermoBrite hybridization system (Abbott Molecular Inc.). The posthybridization wash was with 2X SSC at 73 °C for 3 min followed by a brief water rinse. Slides were air-dried and then counterstained with VECTASHIELD mounting medium with 4’-6-diamidino-2-phenylindole (DAPI) (Vector Laboratories Inc., Burlingame, CA). Image acquisition was performed at 1000x system magnification with a COOL-1300 SpectraCube camera (Applied Spectral Imaging-ASI, Vista, CA) mounted on an Olympus BX43 microscope. Images were analyzed using FISHView v7 software (ASI) and 20 interphase nuclei were scored for each sample.

### Whole exome characterization of uterine leiomyomas and matched normal tissues.

Genomic DNA from all uterine fibroids and corresponding normal myometrium were extracted from freshly frozen tissue using DNeasy Blood and Tissue kit (Qiagen) according to manufactures recommendation. Samples were submitted to Hudson Alpha (Huntsville AL) for 2 x 100 sequencing on an Illumina HiSeq 2500. In total, 8 fibroids and matched normal tissue pairs were sequenced. Three out of the eight sample pairs had an additional fibroid sample sequenced. Exome capture was performed using the NimbleGen SeqCap EZ Exome v3 kit and sequenced to a depth of approximately 45x across two flowcells. Reads were assessed for quality using FastQC v0.11.5 and MultiQC v1.0dev0. Samples were called for germline and somatic variants using the Broad Institute’s “Best Practices” guidelines with GATK v3.6. Briefly, reads were aligned to the human genome (hg19) using BWA mem with the ‐M and ‐R options to mark short, split alignments as secondary and add read group information, respectively. Next, SAM files were converted to coordinate sorted BAM files using samtools v1.3.1, keeping the header (-h) and aligned reads (-F 4). Picard Tools v2.7.1 was used to mark and remove duplicates with the MarkDuplicates functionality and REMOVE_DUPLICATES=true. For germline variant calling, known variant files included: dbSNP build 149, 1000 genomes phase 3, and Mills and 1000 genomes gold standard indels. Interval reference files (design files) for the NimbleGen SeqCap EZ Exome v3 kit were downloaded from the NimbleGen website 04 November, 2016. Specifically, the SeqCap_EZ_Exome_v3_hg19_primary_targets.bed was used with the interval padding option set to 100 bp unless specified otherwise. De-duplicated, aligned reads were subjected to base quality score recalibration with the ‐‐filter_mismatching_base_and_quals and BadCigar filters in place. Calling germline variants was accomplished using HaplotypeCaller with the BadCigar filter in place and emitting a Genomic VCF (GVCF). GVCFs for each sample were combined and genotyped using the GATK GenotypeGVCFs functionality. SNPs were filtered using a QD < 2.0, FS > 60.0, MQ < 40.0, MappingQualityRankSum < −12.5, ReadPosRankSum < −8.0. Indels were filtered using a QD < 2.0, FS > 200.0, ReadPosRankSum < −20.0.

For somatic variant calling, a panel of normals was generated using Mutect2 for each normal sample in artifact detection mode, passing dbSNP, COSMIC (v71), and the NimbleGen SeqCap EZ Exome v3 interval list (SeqCap_EZ_Exome_v3_hg19_primary_targets.bed) with the interval padding option set to 100 bp. For the panel of normals, variants were kept if it was observed in at least two normal samples and not filtered (e.g. ‐‐filteredAreUncalled ‐‐filteredrecordsmergetype KEEP_IF_ANY_UNFILTERED). Variants were called using Mutect2 for each fibroid and normal sample pair passing dbSNP, COSMIC, panel of normals, and the NimbleGen SeqCap EZ Exome v3 interval list to GATK.

Variant annotation was performed on germline and somatic variants using vt (v0.5), VEP (release 87), vcfanno (v0.1.0), and gemini (v0.19.1) in conjunction with COSMIC, ClinVar (Downloaded 12 December 2016), and ExAC databases (Release 0.3.1). First, multi-allelic sites were decomposed and variants normalized using vt. Next, variants were annotated using VEP (Ensembl) and then by vcfanno. A pedigree file (.ped) was generated using the following line: grep ‐m1 CHROM input.vcf | cut ‐f 10‐ | awk ‘BEGIN{RS=“\t”}{ printf(“fam%d\t%s\t0\t0\t-9\t-9\n”, NR, $1)}’ > output.ped. The pedigree file and annotated VCF were then used to generate the gemini database with the vcf2db.py (https://github.com/quinlan-lab/vcf2db). Finally, the annotated variants were filtered and queried against the COSMIC, ClinVar, and ExAC databases using gemini. VCFs were converted to maf using VEP and vcf2maf (https://github.com/mskcc/vcf2maf). Somatic variation was visualized using R (v3.3.2) and maftools (v1.0.55). Whole exome sequencing raw data is made available through SRA (Sequence Read Archive) under SRA identifier SRP163897.

### Illumina Infinium HumanMethylation BeadChip Assay characterization

DNA was quantified by Qubit fluorimetry (Life Technologies) and 500ng of DNA from each sample was bisulfite converted using the Zymo EZ DNA Methylation Kit (Zymo Research, Irvine, CA USA) following the manufacturer’s protocol using the specified modifications for the Illumina Infinium Methylation Assay. After conversion, all bisulfite reactions were cleaned using the Zymo-Spin binding columns, and eluted in 12 uL of Tris buffer. Following elution, BS converted DNA was processed through the EPIC array protocol. To perform the assay, 7uL of converted DNA was denatured with 1ul 0.4N sodium hydroxide. DNA was then amplified, hybridized to the EPIC bead chip, and an extension reaction was performed using fluorophore-labeled nucleotides per the manufacturer’s protocol. Array beadchips were scanned on the Illumina iScan platform.

DNA methylome was captured by the HumanMethylationEPIC (EPIC) BeadChip (Illumina, CA, USA) using the manufacturer’s standard protocol, which interrogates a total of 863,904 CpG loci spreading across the transcription start sites and enhancer/regulatory regions. Additionally, 2932 non-CpG loci (CpH) and 59 single nucleotide polymorphisms (SNPs) are also included as part of the EPIC array.

Raw IDAT files were processed using R package SeSAMe (86) with noob background correction (87), non-linear dye bias correction, and non-detection masking. Methylation beta values were called as quantitative percentage of methylated signals over both unmethylated and methylated signals ranging from 0 to 1, with “0” indicating full lack of methylation and “1” full methylation. We excluded measurements from sub-optimally designed probes due to overlap with SNPs and repeat elements, as suggested by previous studies (16). Raw IDAT files are available through GEO (Gene Expression Omnibus) database under accession GSE120854.

### Methylome analysis

Unsupervised hierarchical clustering was conducted based on most variable probes (top 1% standard deviations out of all the CpG probes) across all samples measured on the EPIC array, and was visualized as heatmap with continuous betas. In order to evaluate the robustness of discovered methylation clusters, we performed consensus clustering by perturbing samples for 1000 iterations (20). Sample‐ and cluster-based stability score were then calculated. Fibroids-specific methylation profiling was generated using CpG probes unmethylated in normal myometria but methylated (defined by a beta value of no less than 0.3) in at least one sample within each fibroid subtype. Hierarchical clustering based on betas within each subtype was then performed to show the specific methylation patterns across different subtypes.

Differentially methylated cytosines (DMCs) were called using R package DMRcate (88) by comparing all the fibroids, and each fibroid subtype to myometria, based on its build-in default p-value cutoff and an absolute beta-value difference threshold at 0.2. DMCs were mapped to genes captured by RNA sequencing, if they were located within gene promoters (defined as 2 kb flanking regions surrounding transcription start sites). We then performed enrichment analysis of transcription factors binding sites (TFBSs) at distal regulatory elements (i.e., enhancers). Distal probes were identified as probes located within TFBSs but not within gene promoters. For hyper‐ or hypo-DMC set generated from each comparison, hypergeometric test was applied to calculate the enrichment or depletion of binding sites for each TF within DMC set at those distal probes. Significance cutoff was made at 1e-6 after false discovery (FDR) correction.

### Evaluating clonality with X inactivation

For patients with more than one fibroids sample examined, we evaluated the possibility that they arose as independent clones or from the same origin. We performed this analysis by integrating exome-seq and RNA-seq data. For each patient, we identified germline SNPs on the X chromosome from exome-seq data using GATK. We restricted the analyses to those SNPs that remained heterozygous in DNA in both tumors. We then examined which alleles (A or B) were expressed for these SNPs using RNA-seq data. Alternative expressed alleles would indicate separate cellular origins, as random inactivation of one X chromosome occurs early in development.

All statistical analyses were conducted using R software with versions newer than 3.4.1 (89).

### Transcriptomic profiling of uterine leiomyomas and matched normal tissues

Total RNA was extracted using Trizol Reagent (Invitrogen) according to manufactures instructions, from freshly frozen samples stored at −80C. The RNA was suspended in RNase-free water, and purified with an RNeasy MinEluteTM clean up kit (Qiagen). RNA concentration and integrity was assessed using a Nanodrop 1000 spectrophotometer (Thermo Scientific, Wilmington, Delaware) and Agilent 2100 Bioanalyzer (Agilent Technologies, Santa Clara, California), according to manufacturer’s protocol. Seventeen samples were submitted to the Van Andel Research Institute (VARI) Genomics Core for 2 x 75 bp RNA sequencing on an Illumina NextSeq 500. Libraries were prepared using a Kapa RNA HyperPrep Kit with ribosomal reduction, pooled, and sequenced across two flowcells to yield approximately 50-60 million reads/sample. Reads were assessed for quality using FastQC v0.11.5 and MulitQC v1.0dev0. Next, raw reads for the sample were merged from two flowcells into a single file and aligned to the human genome (hg19) with STAR v2.5.2b using the two-pass mode. Transcript abundance was quantified using HTSeq v0.6.1p1 with the ‐‐stranded option set to “reverse” and Ensembl GTF (Release 75) as the annotation file. Differential expression (DE) was calculated using either edgeR (v3.16.5) for comparing fibroids to myometria; or limma (v3.30.13) for comparing *HMGA2*hi, and *MED12*mt fibroids to myometria. Counts were filtered to include genes with a minimum of 1 count per million (CPM) in at least 3 samples. Differentially expressed genes were identified as those having a FDR < 0.05 relative to the comparator. MDS plots were generated in R (v3.3.2) using R package ggplot2 (v2.2.1). Expressed somatic variants identified from exomes were determined using the Broad Institute’s “Best Practices” for RNA-seq variant calling. Briefly, we added read group information (using function AddOrReplaceReadGroup from Picard Tools) to BAM files generated by STAR with two-pass mode, and then sorted them by coordinates. Picard Tools v2.7.1 was used to mark duplicates using function MarkDuplicates. Known variant files passed to the exomes were used as known sites in RNA-seq variant calling procedure with GATK v3.6. Cigar strings were modified using function SplitNCigarReads in GATK with the ReassignOneMappingQuality function (RMQF 255, RMQT 60, and ‐U ALLOW_N_CIGAR_READS). Interval targets were generated and indels realigned with GATK. De-duplicated and indel realigned reads were then subjected to base quality score recalibration. After recalibration, these BAMs were fed to HaplotypeCaller to call variants with filters dontUseSoftClippedBases enabled and stand_call_conf set at 20.0. SNPs and indels were further filtered by ‐window 35, ‐cluster 3, FS > 30.0, and QD < 2.0. Expressed somatic variants were identified using bedtools (v2.26.0) with both annotated RNA-seq variants and exome variants. SRA accession number for fastq files is SRP166862.

### Publicly Accessible Data Sets and Bioinformatic Tools

Gene set enrichment analysis was conducted using gene sets downloaded from The Molecular Signatures Database (MSigDB) (90) excluding collection of computational gene sets (C4) and gene ontology gene sets (C5). TFBS-probe annotation of Illumina EPIC array (human reference genome (NCBI build 37/HG19)) was download from (16). Particular gene view was generated using UCSC genome browser with tracks available from track hubs (91), like EPIC probe coordinates, TSS locations, CpG islands, super enhancers (92), enhancer-promoter correlations, DNase peaks, TFBS, Pol2, and CTCF binding signals. The Integrative Genomics Viewer (IGV) was employed to view aligned sequence reads (93).

### Cell culture and nucleofection

UT-TERT myometrial cells were a kind gift from Dr. John Risinger and have been characterized previously (94). UT-TERT cells were cultured and maintained in SmGM™−2 Smooth Muscle Growth Medium-2 containing 5% FBS, 0.1% insulin, 0.2% basic human fibroblast growth factor (hFGF-b), 0.1% GA-100, and 0.1% human epidermal growth factor (hEGF) (Lonza, Walkersville, MD). HOXA13 overexpression plasmid, pLV(Exp)-EGFP:T2A:Puro‐ CBh>hHOXA13, vector ID VB180306-1076naw, was constructed and packaged by VectorBuilder (Cyagen Bioscience). Nucleofection was carried out using Amaxa Basic Nucleofector Kit for Primary Mammalian smooth muscle cells (Lonza, Catalog # VPI-1004) according to manufactures protocol. Briefly, 1×10^6^ cells was resuspended in 100μl Nucleofector solution, and transfected with 1μg of HOXA13 overexpression plasmid using program P-024. Following nucelofection, cells were incubated for 18 hours and media was changed to complete growth media along with supplements. Following 48 hours after nucleofection, the cells were treated with 2μg/ml puromycin to select for stable clones.

### Quantitative Real Time PCR

Total RNA was isolated and treated with Dnase from UT-TERT and HOXA13-UT-TERT clones using an RNA extraction kit (Qiagen, Valencia, CA). cDNA was synthetized from 1 μg of total RNA using SuperScript IV Reverse Transcriptase (Invitrogen). Quantitative Real time PCR (qRT-PCR) analysis using SYBRGreen (BioRad) was performed to analyze gene expression using the ViiA 7 qPCR System (Applied Biosystems). RPL17 was used for normalization. Primer sequences (5’-3’) used for qRT-PCR are *HOXA13* FP (TGGAACGGCCAAATGTACTGCC), *HOXA13* RP (GGTATAAGGCACGCGCTTCTTTC), *DPT* FP (GCCCATATTCCTGCTGGCTAA), DPT RP (GTGGTTGTTGCTCCTCGGAT), *COL3A1* FP (TGGTCTGCAAGGAATGCCTGGA), COL3A1 RP (TCTTTCCCTGGGACACCATCAG), *TGFB3* FP (CTAAGCGGAATGAGCAGAGGATC), *TGFB3* RP (TCTCAACAGCCACTCACGCACA), *HOTTIP* FP (CCTAAAGCCACGCTTCTTTG), *HOTTIP* RP(TGCAGGCTGGAGATCCTACT), *RPL17* FP (ACGAAAAGCCACGAAGTATCTG), *RPL17* RP (GACCTTGTGTCCAGCCCCAT). The fold change in gene expression was calculated using the standard ΔΔCt method.

### Statistical Analysis

Average fibroid and normal tissue samples were calculated within each individual. *HOXA13* and *HOTTIP* expression were then normalized to the housekeeping gene *RPL17*. Kendall’s tau was used to determine the amount of concordance between mean *HOXA13* and *HOTTIP* expression in both fibroid and normal tissues. To determine if mean gene expression normalized to *RPL17* differed between normal and fibroid tissues, a linear mixed-effects model with a random intercept for each patient was fit. These analyses were performed using R v3.4.4 (https://cran.r-project.org) with two-sided hypotheses and a significance level of 0.05.

## REFERENCES

1. Commandeur AE, Styer AK, Teixeira JM. Epidemiological and genetic clues for molecular mechanisms involved in uterine leiomyoma development and growth. Hum Reprod Update 2015; 21:593-615

2. Bulun SE. Uterine fibroids. N Engl J Med 2013; 369:1344-1355

3. Jacoby VL, Fujimoto VY, Giudice LC, Kuppermann M, Washington AE. Racial and ethnic disparities in benign gynecologic conditions and associated surgeries. Am J Obstet Gynecol 2010; 202:514-521

4. Wu JM, Wechter ME, Geller EJ, Nguyen TV, Visco AG. Hysterectomy rates in the United States, 2003. Obstet Gynecol 2007; 110:1091-1095

5. Sandberg AA. Updates on the cytogenetics and molecular genetics of bone and soft tissue tumors: leiomyoma. Cancer Genet Cytogenet 2005; 158:1-26

6. Nilbert M, Heim S, Mandahl N, Floderus UM, Willen H, Mitelman F. Characteristic chromosome abnormalities, including rearrangements of 6p, del(7q), +12, and t(12;14), in 44 uterine leiomyomas. Hum Genet 1990; 85:605-611

7. Zhou X, Benson KF, Przybysz K, Liu J, Hou Y, Cherath L, Chada K. Genomic structure and expression of the murine Hmgi-c gene. Nucleic Acids Res 1996; 24:4071-4077

8. Chen B, Young J, Leng F. DNA bending by the mammalian high-mobility group protein AT hook 2. Biochemistry 2010; 49:1590-1595

9. Makinen N, Mehine M, Tolvanen J, Kaasinen E, Li Y, Lehtonen HJ, Gentile M, Yan J, Enge M, Taipale M, Aavikko M, Katainen R, Virolainen E, Bohling T, Koski TA, Launonen V, Sjoberg J, Taipale J, Vahteristo P, Aaltonen LA. MED12, the mediator complex subunit 12 gene, is mutated at high frequency in uterine leiomyomas. Science 2011; 334:252-255

10. Bertsch E, Qiang W, Zhang Q, Espona-Fiedler M, Druschitz S, Liu Y, Mittal K, Kong B, Kurita T, Wei JJ. MED12 and HMGA2 mutations: two independent genetic events in uterine leiomyoma and leiomyosarcoma. Mod Pathol 2014; 27:1144-1153

11. Mehine M, Makinen N, Heinonen HR, Aaltonen LA, Vahteristo P. Genomics of uterine leiomyomas: insights from high-throughput sequencing. Fertil Steril 2014; 102:621-629

12. Jones PA. Functions of DNA methylation: islands, start sites, gene bodies and beyond. Nat Rev Genet 2012; 13:484-492

13. Mehine M, Kaasinen E, Heinonen HR, Makinen N, Kampjarvi K, Sarvilinna N, Aavikko M, Vaharautio A, Pasanen A, Butzow R, Heikinheimo O, Sjoberg J, Pitkanen E, Vahteristo P, Aaltonen LA. Integrated data analysis reveals uterine leiomyoma subtypes with distinct driver pathways and biomarkers. Proc Natl Acad Sci U S A 2016; 113:1315-1320

14. Lister R, Mukamel EA, Nery JR, Urich M, Puddifoot CA, Johnson ND, Lucero J, Huang Y, Dwork AJ, Schultz MD, Yu M, Tonti-Filippini J, Heyn H, Hu S, Wu JC, Rao A, Esteller M, He C, Haghighi FG, Sejnowski TJ, Behrens MM, Ecker JR. Global epigenomic reconfiguration during mammalian brain development. Science 2013; 341:1237905

15. Lister R, Pelizzola M, Dowen RH, Hawkins RD, Hon G, Tonti-Filippini J, Nery JR, Lee L, Ye Z, Ngo QM, Edsall L, Antosiewicz-Bourget J, Stewart R, Ruotti V, Millar AH, Thomson JA, Ren B, Ecker JR. Human DNA methylomes at base resolution show widespread epigenomic differences. Nature 2009; 462:315-322

16. Zhou W, Laird PW, Shen H. Comprehensive characterization, annotation and innovative use of Infinium DNA methylation BeadChip probes. Nucleic Acids Res 2017; 45:e22

17. Stewart EA, Nicholson WK, Bradley L, Borah BJ. The burden of uterine fibroids for African-American women: results of a national survey. J Womens Health (Larchmt) 2013; 22:807-816

18. Mongroo PS, Rustgi AK. The role of the miR-200 family in epithelial-mesenchymal transition. Cancer Biol Ther 2010; 10:219-222

19. Hu B, Gharaee-Kermani M, Wu Z, Phan SH. Epigenetic regulation of myofibroblast differentiation by DNA methylation. Am J Pathol 2010; 177:21-28

20. Wilkerson MD, Hayes DN. ConsensusClusterPlus: a class discovery tool with confidence assessments and item tracking. Bioinformatics 2010; 26:1572-1573

21. Chalmers ZR, Connelly CF, Fabrizio D, Gay L, Ali SM, Ennis R, Schrock A, Campbell B, Shlien A, Chmielecki J, Huang F, He Y, Sun J, Tabori U, Kennedy M, Lieber DS, Roels S, White J, Otto GA, Ross JS, Garraway L, Miller VA, Stephens PJ, Frampton GM. Analysis of 100,000 human cancer genomes reveals the landscape of tumor mutational burden. Genome Med 2017; 9:34

22. Zhou W, Dinh HQ, Ramjan Z, Weisenberger DJ, Nicolet CM, Shen H, Laird PW, Berman BP. DNA methylation loss in late-replicating domains is linked to mitotic cell division. Nat Genet 2018; 50:591-602

23. Yao L, Shen H, Laird PW, Farnham PJ, Berman BP. Inferring regulatory element landscapes and transcription factor networks from cancer methylomes. Genome Biol 2015; 16:105

24. Consortium EP. An integrated encyclopedia of DNA elements in the human genome. Nature 2012; 489:57-74

25. Margueron R, Reinberg D. The Polycomb complex PRC2 and its mark in life. Nature 2011; 469:343-349

26. Ashar HR, Fejzo MS, Tkachenko A, Zhou X, Fletcher JA, Weremowicz S, Morton CC, Chada K. Disruption of the architectural factor HMGI-C: DNA-binding AT hook motifs fused in lipomas to distinct transcriptional regulatory domains. Cell 1995; 82:57-65

27. Schoenmakers EF, Wanschura S, Mols R, Bullerdiek J, Van den Berghe H, Van de Ven WJ. Recurrent rearrangements in the high mobility group protein gene, HMGI-C, in benign mesenchymal tumours. Nat Genet 1995; 10:436-444

28. Wang J, Zhuang J, Iyer S, Lin X, Whitfield TW, Greven MC, Pierce BG, Dong X, Kundaje A, Cheng Y, Rando OJ, Birney E, Myers RM, Noble WS, Snyder M, Weng Z. Sequence features and chromatin structure around the genomic regions bound by 119 human transcription factors. Genome Res 2012; 22:1798-1812

29. Wang H, Maurano MT, Qu H, Varley KE, Gertz J, Pauli F, Lee K, Canfield T, Weaver M, Sandstrom R, Thurman RE, Kaul R, Myers RM, Stamatoyannopoulos JA. Widespread plasticity in CTCF occupancy linked to DNA methylation. Genome Res 2012; 22:1680-1688

30. Fortin JP, Hansen KD. Reconstructing A/B compartments as revealed by Hi-C using long-range correlations in epigenetic data. Genome Biol 2015; 16:180

31. Navarro A, Yin P, Monsivais D, Lin SM, Du P, Wei JJ, Bulun SE. Genome-wide DNA methylation indicates silencing of tumor suppressor genes in uterine leiomyoma. PLoS One 2012; 7:e33284

32. Casey R, Rogers PA, Vollenhoven BJ. An immunohistochemical analysis of fibroid vasculature. Human reproduction 2000; 15:1469-1475

33. Mayer A, Hockel M, Wree A, Leo C, Horn LC, Vaupel P. Lack of hypoxic response in uterine leiomyomas despite severe tissue hypoxia. Cancer Res 2008; 68:4719-4726

34. Taylor HS, Vanden Heuvel GB, Igarashi P. A conserved Hox axis in the mouse and human female reproductive system: late establishment and persistent adult expression of the Hoxa cluster genes. Biol Reprod 1997; 57:1338-1345

35. Warot X, Fromental-Ramain C, Fraulob V, Chambon P, Dolle P. Gene dosage-dependent effects of the Hoxa-13 and Hoxd-13 mutations on morphogenesis of the terminal parts of the digestive and urogenital tracts. Development 1997; 124:4781-4791

36. Zhao Y, Potter SS. Functional specificity of the Hoxa13 homeobox. Development 2001; 128:3197-3207

37. Scott V, Morgan EA, Stadler HS. Genitourinary functions of Hoxa13 and Hoxd13. J Biochem 2005; 137:671-676

38. Daftary GS, Taylor HS. Endocrine regulation of HOX genes. Endocr Rev 2006; 27:331-355

39. Wang KC, Yang YW, Liu B, Sanyal A, Corces-Zimmerman R, Chen Y, Lajoie BR, Protacio A, Flynn RA, Gupta RA, Wysocka J, Lei M, Dekker J, Helms JA, Chang HY. A long noncoding RNA maintains active chromatin to coordinate homeotic gene expression. Nature 2011; 472:120-124

40. Malik M, Norian J, McCarthy-Keith D, Britten J, Catherino WH. Why leiomyomas are called fibroids: the central role of extracellular matrix in symptomatic women. Semin Reprod Med 2010; 28:169-179

41. Arslan AA, Gold LI, Mittal K, Suen TC, Belitskaya-Levy I, Tang MS, Toniolo P. Gene expression studies provide clues to the pathogenesis of uterine leiomyoma: new evidence and a systematic review. Human reproduction 2005; 20:852-863

42. Kobayashi A, Behringer RR. Developmental genetics of the female reproductive tract in mammals. Nat Rev Genet 2003; 4:969-980

43. Sun NX, Ye C, Zhao Q, Zhang Q, Xu C, Wang SB, Jin ZJ, Sun SH, Wang F, Li W. Long noncoding RNA-EBIC promotes tumor cell invasion by binding to EZH2 and repressing E-cadherin in cervical cancer. PLoS One 2014; 9:e100340

44. Pappa KI, Polyzos A, Jacob-Hirsch J, Amariglio N, Vlachos GD, Loutradis D, Anagnou NP. Profiling of Discrete Gynecological Cancers Reveals Novel Transcriptional Modules and Common Features Shared by Other Cancer Types and Embryonic Stem Cells. PLoS One 2015; 10:e0142229

45. Mittal P, Shin YH, Yatsenko SA, Castro CA, Surti U, Rajkovic A. Med12 gain-of-function mutation causes leiomyomas and genomic instability. J Clin Invest 2015; 125:3280-3284

46. Park MJ, Shen H, Spaeth JM, Tolvanen JH, Failor C, Knudtson JF, McLaughlin J, Halder SK, Yang Q, Bulun SE, Al-Hendy A, Schenken RS, Aaltonen LA, Boyer TG. Oncogenic exon 2 mutations in Mediator subunit MED12 disrupt allosteric activation of cyclin CCDK8/19. J Biol Chem 2018; 293:4870-4882

47. Quade BJ, Weremowicz S, Neskey DM, Vanni R, Ladd C, Dal Cin P, Morton CC. Fusion transcripts involving HMGA2 are not a common molecular mechanism in uterine leiomyomata with rearrangements in 12q15. Cancer Res 2003; 63:1351-1358

48. Schoenberg Fejzo M, Ashar HR, Krauter KS, Powell WL, Rein MS, Weremowicz S, Yoon SJ, Kucherlapati RS, Chada K, Morton CC. Translocation breakpoints upstream of the HMGIC gene in uterine leiomyomata suggest dysregulation of this gene by a mechanism different from that in lipomas. Genes Chromosomes Cancer 1996; 17:1-6

49. Peng Y, Laser J, Shi G, Mittal K, Melamed J, Lee P, Wei JJ. Antiproliferative effects by Let-7 repression of high-mobility group A2 in uterine leiomyoma. Mol Cancer Res 2008; 6:663-673

50. Wang T, Zhang X, Obijuru L, Laser J, Aris V, Lee P, Mittal K, Soteropoulos P, Wei JJ. A micro-RNA signature associated with race, tumor size, and target gene activity in human uterine leiomyomas. Genes Chromosomes Cancer 2007; 46:336-347

51. Takahashi T, Nagai N, Oda H, Ohama K, Kamada N, Miyagawa K. Evidence for RAD51L1/HMGIC fusion in the pathogenesis of uterine leiomyoma. Genes Chromosomes Cancer 2001; 30:196-201

52. Phillips JE, Corces VG. CTCF: master weaver of the genome. Cell 2009; 137:1194-1211

53. Calo E, Wysocka J. Modification of enhancer chromatin: what, how, and why? Mol Cell 2013; 49:825-837

54. Shlyueva D, Stampfel G, Stark A. Transcriptional enhancers: from properties to genome-wide predictions. Nat Rev Genet 2014; 15:272-286

55. Sur I, Taipale J. The role of enhancers in cancer. Nat Rev Cancer 2016; 16:483-493

56. Maekawa R, Sato S, Yamagata Y, Asada H, Tamura I, Lee L, Okada M, Tamura H, Takaki E, Nakai A, Sugino N. Genome-wide DNA methylation analysis reveals a potential mechanism for the pathogenesis and development of uterine leiomyomas. PLoS One 2013; 8:e66632

57. Simon JA, Kingston RE. Occupying chromatin: Polycomb mechanisms for getting to genomic targets, stopping transcriptional traffic, and staying put. Mol Cell 2013; 49:808-824

58. Widschwendter M, Fiegl H, Egle D, Mueller-Holzner E, Spizzo G, Marth C, Weisenberger DJ, Campan M, Young J, Jacobs I, Laird PW. Epigenetic stem cell signature in cancer. Nat Genet 2007; 39:157-158

59. Yang Q, Nair S, Laknaur A, Ismail N, Diamond MP, Al-Hendy A. The Polycomb Group Protein EZH2 Impairs DNA Damage Repair Gene Expression in Human Uterine Fibroids. Biol Reprod 2016; 94:69

60. Boyer LA, Lee TI, Cole MF, Johnstone SE, Levine SS, Zucker JP, Guenther MG, Kumar RM, Murray HL, Jenner RG, Gifford DK, Melton DA, Jaenisch R, Young RA. Core transcriptional regulatory circuitry in human embryonic stem cells. Cell 2005; 122:947-956

61. Ono M, Qiang W, Serna VA, Yin P, Coon JSt, Navarro A, Monsivais D, Kakinuma T, Dyson M, Druschitz S, Unno K, Kurita T, Bulun SE. Role of stem cells in human uterine leiomyoma growth. PLoS One 2012; 7:e36935

62. Chang HL, Senaratne TN, Zhang L, Szotek PP, Stewart E, Dombkowski D, Preffer F, Donahoe PK, Teixeira J. Uterine leiomyomas exhibit fewer stem/progenitor cell characteristics when compared with corresponding normal myometrium. Reprod Sci 2010; 17:158-167

63. Benassayag C, Leroy MJ, Rigourd V, Robert B, Honore JC, Mignot TM, Vacher-Lavenu MC, Chapron C, Ferre F. Estrogen receptors (ERalpha/ERbeta) in normal and pathological growth of the human myometrium: pregnancy and leiomyoma. Am J Physiol 1999; 276:E1112-1118

64. Kovacs KA, Oszter A, Gocze PM, Kornyei JL, Szabo I. Comparative analysis of cyclin D1 and oestrogen receptor (alpha and beta) levels in human leiomyoma and adjacent myometrium. Mol Hum Reprod 2001; 7:1085-1091

65. Otsuka H, Shinohara M, Kashimura M, Yoshida K, Okamura Y. A comparative study of the estrogen receptor ratio in myometrium and uterine leiomyomas. Int J Gynaecol Obstet 1989; 29:189-194

66. Barbarisi A, Petillo O, Di Lieto A, Melone MA, Margarucci S, Cannas M, Peluso G. 17-beta estradiol elicits an autocrine leiomyoma cell proliferation: evidence for a stimulation of protein kinase-dependent pathway. J Cell Physiol 2001; 186:414-424

67. Bever AT, Hisaw FL, Velardo JT. Inhibitory action of desoxycorticosterone acetate, cortisone acetate, and testosterone on uterine growth induced by estradiol-17beta. Endocrinology 1956; 59:165-169

68. Rhen T, Grissom S, Afshari C, Cidlowski JA. Dexamethasone blocks the rapid biological effects of 17beta-estradiol in the rat uterus without antagonizing its global genomic actions. FASEB J 2003; 17:1849-1870

69. Bitman J, Cecil HC. Differential inhibition by cortisol of estrogen-stimulated uterine responses. Endocrinology 1967; 80:423-429

70. Hennig Y, Deichert U, Bonk U, Thode B, Bartnitzke S, Bullerdiek J. Chromosomal translocations affecting 12q14-15 but not deletions of the long arm of chromosome 7 associated with a growth advantage of uterine smooth muscle cells. Mol Hum Reprod 1999; 5:1150-1154

71. Heinonen HR, Pasanen A, Heikinheimo O, Tanskanen T, Palin K, Tolvanen J, Vahteristo P, Sjoberg J, Pitkanen E, Butzow R, Makinen N, Aaltonen LA. Multiple clinical characteristics separate MED12-mutation-positive and ‐negative uterine leiomyomas. Sci Rep 2017; 7:1015

72. Chang S, Liu J, Guo S, He S, Qiu G, Lu J, Wang J, Fan L, Zhao W, Che X. HOTTIP and HOXA13 are oncogenes associated with gastric cancer progression. Oncol Rep 2016; 35:3577-3585

73. Duan R, Han L, Wang Q, Wei J, Chen L, Zhang J, Kang C, Wang L. HOXA13 is a potential GBM diagnostic marker and promotes glioma invasion by activating the Wnt and TGF-beta pathways. Oncotarget 2015; 6:27778-27793

74. Luo Z, Rhie SK, Lay FD, Farnham PJ. A Prostate Cancer Risk Element Functions as a Repressive Loop that Regulates HOXA13. Cell Rep 2017; 21:1411-1417

75. Yamashita T, Tazawa S, Yawei Z, Katayama H, Kato Y, Nishiwaki K, Yokohama Y, Ishikawa M. Suppression of invasive characteristics by antisense introduction of overexpressed HOX genes in ovarian cancer cells. Int J Oncol 2006; 28:931-938

76. Larsen B, Hwang J. Progesterone interactions with the cervix: translational implications for term and preterm birth. Infect Dis Obstet Gynecol 2011; 2011:353297

77. Ishikawa H, Ishi K, Serna VA, Kakazu R, Bulun SE, Kurita T. Progesterone is essential for maintenance and growth of uterine leiomyoma. Endocrinology 2010; 151:2433-2442

78. Timmons B, Akins M, Mahendroo M. Cervical remodeling during pregnancy and parturition. Trends Endocrinol Metab 2010; 21:353-361

79. Baird DD, Dunson DB. Why is parity protective for uterine fibroids? Epidemiology 2003; 14:247-250

80. Vodstrcil LA, Shynlova O, Westcott K, Laker R, Simpson E, Wlodek ME, Parry LJ. Progesterone withdrawal, and not increased circulating relaxin, mediates the decrease in myometrial relaxin receptor (RXFP1) expression in late gestation in rats. Biol Reprod 2010; 83:825-832

81. Challis JRG, Matthews SG, Gibb W, Lye SJ. Endocrine and paracrine regulation of birth at term and preterm. Endocr Rev 2000; 21:514-550

82. Liu L, Li H, Dargahi D, Shynlova O, Slater D, Jones SJ, Lye SJ, Dong X. HoxA13 Regulates Phenotype Regionalization of Human Pregnant Myometrium. J Clin Endocrinol Metab 2015; 100:E1512-1522

83. Park SM, Gaur AB, Lengyel E, Peter ME. The miR-200 family determines the epithelial phenotype of cancer cells by targeting the E-cadherin repressors ZEB1 and ZEB2. Genes Dev 2008; 22:894-907

84. Vrba L, Jensen TJ, Garbe JC, Heimark RL, Cress AE, Dickinson S, Stampfer MR, Futscher BW. Role for DNA methylation in the regulation of miR-200c and miR-141 expression in normal and cancer cells. PLoS One 2010; 5:e8697

85. Skalli O, Schurch W, Seemayer T, Lagace R, Montandon D, Pittet B, Gabbiani G. Myofibroblasts from diverse pathologic settings are heterogeneous in their content of actin isoforms and intermediate filament proteins. Lab Invest 1989; 60:275-285

86. Zhou W, Triche TJ, Jr., Laird PW, Shen H. SeSAMe: reducing artifactual detection of DNA methylation by Infinium BeadChips in genomic deletions. Nucleic Acids Res 2018;

87. Triche TJ, Jr., Weisenberger DJ, Van Den Berg D, Laird PW, Siegmund KD. Low-level processing of Illumina Infinium DNA Methylation BeadArrays. Nucleic Acids Res 2013; 41:e90

88. Peters TJ, Buckley MJ, Statham AL, Pidsley R, Samaras K, R VL, Clark SJ, Molloy PL. De novo identification of differentially methylated regions in the human genome. Epigenetics Chromatin 2015; 8:6

89. R: A Language and Environment for Statistical Computing [computer program]. 2006 Vienna, Austria: R Foundation for Statistical Computing.

90. Subramanian A, Tamayo P, Mootha VK, Mukherjee S, Ebert BL, Gillette MA, Paulovich A, Pomeroy SL, Golub TR, Lander ES, Mesirov JP. Gene set enrichment analysis: a knowledge-based approach for interpreting genome-wide expression profiles. Proc Natl Acad Sci U S A 2005; 102:15545-15550

91. Kent WJ, Sugnet CW, Furey TS, Roskin KM, Pringle TH, Zahler AM, Haussler D. The human genome browser at UCSC. Genome Res 2002; 12:996-1006

92. Wei Y, Zhang S, Shang S, Zhang B, Li S, Wang X, Wang F, Su J, Wu Q, Liu H, Zhang Y. SEA: a super-enhancer archive. Nucleic Acids Res 2016; 44:D172-179

93. Robinson JT, Thorvaldsdottir H, Winckler W, Guttman M, Lander ES, Getz G, Mesirov JP. Integrative genomics viewer. Nat Biotechnol 2011; 29:24-26

94. Carney SA, Tahara H, Swartz CD, Risinger JI, He H, Moore AB, Haseman JK, Barrett JC, Dixon D. Immortalization of human uterine leiomyoma and myometrial cell lines after induction of telomerase activity: molecular and phenotypic characteristics. Lab Invest 2002; 82:719-728

